# Maternal cardiometabolic dysfunction and fetal sex-specific alterations to uterine vascular reactivity in an ovine model of diet-induced obesity during pregnancy

**DOI:** 10.64898/2026.01.15.699637

**Authors:** Rachael C. Crew, Anna L.K. Cochrane, Youguo Niu, Sage G. Ford, Clement L.R. Cahen, Skaai H. Davison, Michael P. Murphy, Susan E. Ozanne, Dino A. Giussani

## Abstract

Obesity during pregnancy is at pandemic proportions and predisposes women to pre- and postnatal cardiovascular dysfunction. The mechanisms underlying this maternal cardiovascular vulnerability remain unclear, partly due to a lack of translatable models capable of longitudinal *in vivo* cardiovascular monitoring. Here, we characterise a novel ovine model of maternal diet-induced obesity during pregnancy. Ewes were fed a control (CON) or obesogenic (OB; *ad libitum* concentrates) diet for 60 days pre-pregnancy and throughout gestation. Pregnant ewes were surgically instrumented with vascular catheters and Transonic flow probes using the wireless CamDAS system, which measured maternal cardiovascular function near term in free-moving ewes. Uterine artery vasoreactivity was assessed *ex vivo* by *in vitro* wire myography. OB ewes entered pregnancy 30% heavier than controls (*P*<0.003) and were hyperglycaemic, hyperinsulinemic, and hyperlipidaemic during pregnancy, relative to CON ewes (all *P*<0.05). OB ewes had elevated haematocrit and haemoglobin across pregnancy, and were hypertensive near term, with an increase in basal femoral artery blood flow, and elevated peripheral oxygen and glucose delivery (all *P*<0.05). OB mothers carrying a female fetus showed increased uterine artery vascular resistance *in vivo* (*P*<0.005) and reduced smooth muscle-dependent vasorelaxation *ex vivo* (*P*<0.05) relative to CON. Conversely, OB mothers carrying a male fetus showed greater NO-independent mechanisms mediating the uterine vasodilator response to methacholine *ex vivo* (*P*<0.001). Collectively, this study characterises a robust model of maternal obesity during pregnancy that offers clinical translational potential and highlights fetal sex-specific changes to uterine artery function.

**Key points summary:**

- Obesity during pregnancy is increasingly common and predisposes women to cardiovascular dysfunction during pregnancy and long after birth, but the specific mechanisms underlying this remain unclear.
- We developed a novel ovine model of diet-induced obesity during pregnancy that displays maternal hypertension, elevated haemoglobin, metabolic dysfunction and alterations in uterine and peripheral blood flow and nutrient delivery near term.
- Mothers with obesity carrying a female fetus had elevated uterine vascular resistance *in vivo* and reduced uterine artery smooth muscle-dependent vasodilator reactivity *ex vivo*.
- Mothers with obesity carrying a male fetus showed no effect on uterine vascular resistance *in vivo*, but greater NO-independent mechanisms mediating the uterine vasodilator response to methacholine *ex vivo*.
- These findings highlight that fetal sex may influence maternal cardiovascular function during obese pregnancy.

## Introduction

Women continue to be underdiagnosed, undertreated, and underrepresented in cardiovascular science, with research failing to sufficiently address factors that uniquely affect a woman’s cardiovascular risk across the life course (Vervoort *et al*., 2024). Women who develop gestational complications are more susceptible to cardiovascular dysfunction during pregnancy, and this heightened cardiovascular vulnerability can continue long after delivery (Parikh *et al*., 2021; Täufer Cederlöf *et al*., 2022). Maternal obesity is a major risk factor for pregnancy complications, which is of the gravest concern given that rates of obesity in women of reproductive age have reached pandemic proportions (Kent *et al*., 2024; Schon *et al*., 2024). However, the mechanistic links between obesity during pregnancy and maternal cardiovascular dysfunction remain unclear, partly due to a lack of experiments in animal models of increased human translational potential that permit invasive longitudinal *in vivo* monitoring of maternal cardiometabolic function.

The ovine model offers distinct translational advantages in pregnancy research, thereby bridging the gap between human populations and mechanistic preclinical studies. Ovine studies add a new dimension to what can be addressed by the widely used rodent models of maternal obesity during pregnancy, which are used by many laboratories, including ours (see Cochrane *et al*. (2024) for review). In contrast to altricial, litter bearing rodents, sheep share similar developmental milestones to humans, exhibiting a relatively long gestational period, and giving birth to precocial singleton or twin offspring with comparable birth weights to humans (Morrison *et al*., 2018). The ovine placenta, while anatomically distinct, shares physiological similarities with humans, including counter-current flow of maternal and fetal blood within the placental villous tree, almost identical oxygen gradients and consumption rates (37 versus 34 mL·kg^-1^·min^-1^ in sheep and humans, respectively), and similar glucose transfer mechanisms and nutrient transporter expression profiles (Bonds *et al*., 1986; Wilkening *et al*., 1988; Barry & Anthony, 2008; Ma *et al*., 2011; Regnault *et al*., 2013). These similarities in fetal size and nutritional demand induce comparable maternal cardiometabolic adaptations to pregnancy between sheep and humans. Moreover, surgical instrumentation of the maternal-fetal vasculature is possible in sheep, which enables serial blood sampling and longitudinal measurement of *in vivo* cardiometabolic function (Allison *et al*., 2016; Tong *et al*., 2022).

Here, we report the development and characterisation of a novel ovine model of obesity during pregnancy that displays maternal metabolic dysfunction, hypertension, and alterations in uterine and peripheral blood flow and nutrient delivery near term. We further show fetal sex-specific alterations to *in vivo* uterine artery function and *ex vivo* uterine artery reactivity in mothers with obesity, highlighting the impact of maternal-fetal communication on maternal vascular adaptations to pregnancy. This offers mechanistic insight into vascular disruption in a clinically accessible and routinely monitored vascular bed, thereby providing enhanced translational potential.

## Materials and Methods

### Ethical approval

These studies were conducted in multiparous 2–3-year-old Welsh Mountain ewes at the Barcroft Centre of the University of Cambridge. All procedures involving animals were performed under the UK Animals (Scientific Procedures) Act 1986 (Project licences: PC6CEFE59/PP6755721), following approval by the University of Cambridge Animal Welfare and Ethical Review Board. All investigators adhered to the ethical principles and reporting standards for animal experiments outlined by Grundy *et al*. (2015). The experimental design followed the recommendations of the ARRIVE (2012) and the National Centre for Replacement Refinement and Reduction (NC3Rs) guidelines (Tannenbaum & Bennett, 2015).

### Feeding regimen and pregnancy establishment

Following a minimum of two weeks of acclimatisation to the facility, Welsh Mountain ewes were fed a control diet (CON, recommended ration, 200g/day/sheep, Bearts Ewe Nuts; H & C Beart Ltd, Norfolk, UK, and *ad libitum* hay), or an obesogenic diet (OB, *ad libitum* access to concentrates and hay) for at least 60 days pre-pregnancy and throughout gestation. Body condition scores between 1 (emaciated) and 5 (obese) were obtained bi-weekly by two of all trained assessors (RCC, SGF, YN, ALKC, CLRC) via palpation of the transverse and vertical processes of the lumbar vertebrae (Kenyon *et al*., 2014). Following 8-10 weeks of the feeding regimen, oestrus was synchronised in ewes via the insertion of a controlled release flugestone acetate vaginal sponge (Chronogest® CR 20 mg). Ewes were then housed with a stud ram for five days, with the date of raddle marking designated as day 0 of gestation. Pregnancy was confirmed by an ultrasound scan at 80 days of gestational age (dGA). Term in this breed is *ca.* 147 days (Brain *et al*., 2019). All ewes were maintained on their respective CON or OB diet throughout the experimental period.

### Longitudinal blood sampling protocol

Serial blood samples (10 ml) were taken from the external jugular vein in a subset of ewes at baseline (prior to CON or OB diet allocation), after four and eight weeks of pre-pregnancy diet exposure, at marking (peri-conception), and at 40, 80 and 120 dGA. Blood samples were analysed for glucose, haemoglobin (Hb) and haematocrit (Hct) with an ABL90 FLEX PLUS blood gas analyser (Radiometer Ltd, Crawley, UK). The remaining EDTA-prepared blood samples were centrifuged at 2370 x *g* for 5 min. Plasma aliquots were snap-frozen in liquid nitrogen and stored at -80°C until further analyses.

### Maternal surgical instrumentation

Under general anaesthesia, a subset of pregnant ewes (*n*= 8 CON, 12 OB) were surgically instrumented with vascular catheters and perivascular flow probes to measure maternal cardiovascular function, as described previously (Allison *et al*., 2016; Allison *et al*., 2020; Tong *et al*., 2022). Briefly, ewes were fasted for 24 h prior to surgery with continuous access to water. On the day of surgery, anaesthesia was induced in animals with a jugular vein injection of Alfaxan (1.5-2.5 mg/kg alfaxalone; Jurox Ltd., UK). Ewes were then intubated (Portex cuffed endotracheal tube; Smiths Medical International Ltd., UK) using a laryngoscope for maintenance of general anaesthesia using 1.5-2.0% isoflurane (IsoFlo; Abbott Laboratories Ltd., UK) in 60:40 O_2_:N_2_O during spontaneous breathing. The maternal abdomen, flanks and medial surfaces of the hind limbs were then shaved and cleaned, and pre-operative antibiotics (30 mg/kg procaine benzylpenicillin i.m.; Depocillin; Intervet UK Ltd., UK) and an analgesic agent (1.4 mg/kg carprofen s.c.; Rimadyl; Pfizer Ltd., UK) were administered. The ewe was then transferred to the surgical theatre and general anaesthesia was maintained using a positive pressure ventilator (Datex-Ohmeda Ltd., UK). The animal was covered with sterile drapes, a midline abdominal incision was made and a Transonic flow probe (MC4PSB-JS-WX120-CM4B-GC; Transonic Systems Europe, Netherlands) was positioned around one of the main uterine arteries. Bilateral incisions were made within the maternal femoral triangles, and catheters were inserted into the right maternal femoral artery (Teflon; ID, 1.0 mm; OD, 1.6 mm; Altec, UK) and vein (polyvinyl chloride; ID, 0.86 mm; OD, 1.52 mm; Critchley Electrical Products, NSW, Australia) with the tips placed in the descending aorta and inferior vena cava, respectively. A second Transonic flow (MC4PSB-JS-WX120-CM4B-GC) probe was implanted around the left maternal femoral artery. All catheters and flow probe leads were exteriorized through keyhole incisions on the ewe’s flanks. While under general anaesthesia, the ewe was fitted with a bespoke jacket housing the CamDAS, a wireless data acquisition system developed in our laboratory (Allison *et al*., 2016; Allison *et al*., 2020; Tong *et al*., 2022). The arterial catheter was connected to a pressure transducer within the pressure box unit, and the Transonic flow probes were connected to the flow box unit housed in the jacket (Allison *et al*., 2016; Allison *et al*., 2020; Tong *et al*., 2022). The dead space of each catheter was then filled with heparinized saline (100 IU/ml heparin in 0.9% NaCl). Isoflurane was withdrawn and the ewe was extubated after spontaneous breathing had returned.

During post-operative recovery, ewes were housed in individual floor pens under a 12:12h light:dark cycle and maintained on their respective CON or OB diets. Analgesia and antibiotic administration continued for 3-5 days following surgery, as described previously (Allison *et al*., 2020). All catheters were flushed daily with heparinized saline to maintain patency, and maternal arterial blood gas measurements were taken daily to monitor ewe wellbeing.

### Maternal cardiovascular recording and nutrient delivery calculations

After five days of post-operative recovery, descending aortic blood pressure (via the femoral artery catheter) and blood flow in the femoral and uterine arteries were recorded continuously on a beat-to-beat basis in each free-moving ewe. Cardiovascular data were visualised on IDEEQ 2.15.0 software and analysed in LabChart 8 to calculate the systolic, diastolic, and mean arterial blood pressure. Heart rate was calculated from the femoral artery flow pulse. Vascular resistance in the uterine and femoral arterial circulations was calculated using Ohm’s principle, by dividing the mean arterial blood pressure by the respective blood flow. Blood gas samples were taken from the femoral artery catheter at the start of the cardiovascular recording period. The maternal arterial blood oxygen content (C_a_O_2_) was then calculated as:

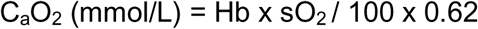

where Hb is haemoglobin concentration (g/L); sO_2_ is oxygen saturation (%), and one molecule of Hb (MW 64.450) binds four molecules of oxygen, as described previously (Gardner *et al*., 2003; Allison *et al*., 2016; Allison *et al*., 2020). The contribution of oxygen dissolved in plasma was considered negligible (Owens *et al*., 1987).

Oxygen and glucose delivery to the maternal peripheral and uterine vascular beds was then calculated according to the following equations:

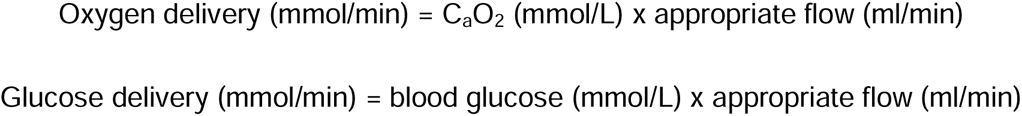

### Post-mortem tissue collection

At 130 dGA, blood samples were collected from the maternal jugular vein in a subset of non-instrumented animals (*n*=18 CON, 18 OB). Maternal blood pH, partial pressure of oxygen (PO_2_) and carbon dioxide (pCO_2_), oxygen saturation (sO_2_), haemoglobin (Hb), glucose, lactate and electrolytes were measured with the ABL90 FLEX PLUS blood gas analyser (Radiometer Ltd, Crawley, UK). Ewes in the instrumented and non-instrumented groups were then humanely killed by an overdose of sodium pentobarbitone (0.4 ml/kg i.v Pentoject; Animal Ltd, York, United Kingdom). The fetus was exteriorised via Caesarean section and weighed. Fetal measurements were taken, including the crown rump length (CRL), biparietal diameter (BPD), abdominal circumference (AC) and femur length. Fetal BMI was calculated as Fetal Body Weight (kg) / CRL (cm)^2^ and Ponderal Index (PI) was calculated as Fetal Body Weight (g) x 100 / CRL (cm)^3^. Placentomes were isolated and classified as A, B, C or D subtypes, according to Vatnick et al. (1991). Each placentome type was counted and weighed individually. Maternal and fetal organs were also isolated and weighed. For bilateral organs, the right maternal organ and both fetal organs were weighed. When bilateral fetal organs showed no statistical difference according to anatomical position, weights were presented as the mean of both organs.

### Uterine artery wire myography

Following post-mortem at 130 dGA, uterine artery reactivity was determined in a subset of ewes (*n*=12 CON, 13 OB) by *in vitro* wire myography. A third-order branch of the uterine artery was isolated, and *ca.* 2 mm vessel segments were threaded with stainless steel 40 µm diameter wire and secured in microvascular chambers (Wire Myograph System 610M; DMT, Aarhus, Denmark). The optimal diameter was determined for each vessel according to Delaey et al. (2002). Briefly, vessels were stretched to a diameter of 400 μm and equilibrated in Krebs buffer (118.5 mM NaCl, 25 mM NaHCO_3_, 4.7 mM KCl, 1.2 mM MgSO_4_.7H2O, 1.2 mM KH_2_PO_4_, 2.5 mM CaCl_2_, 2.8 mM d-glucose; Sigma-Aldrich, Gillingham, UK) bubbled with 95% O_2_/5% CO_2_ at 37°C. Once the tension recording was stable, the chamber was filled with High K^+^ Krebs (59.25 mM NaCl, 25 mM NaHCO_3_, 4.7 mM KCl, 1.2 mM MgSO_4_.7H_2_O, 64.86 mM KH_2_PO_4_, 2.5 mM CaCl_2_, 2.8 mM d-glucose; Sigma-Aldrich, Gillingham, UK) and the maximum change in vessel tension recorded. Vessels were then washed in standard Krebs buffer and the vessel diameter increased by 25-100 μm increments, depending on the magnitude of response. When tension returned to baseline, the high K^+^ Krebs was again administered and the maximum change in tension recorded. These steps were repeated until the change in tension was equal to that reached at the previous vessel diameter, indicating that an optimal physiological working diameter was achieved (Delaey *et al*., 2002). Vessels were then washed with standard Krebs buffer and left to rest for at least 20 min before the generation of dose-response curves.

Uterine vascular constrictor function was determined by measuring tension to cumulative increasing doses of serotonin (5-HT; 10^-9^ to 10^-4^ M). This 5-HT dose-response curve was then repeated following a 20 min pre-incubation with the synthetic Rho-kinase inhibitor Y27632 to determine the Rho kinase-dependent contribution to the vasoconstriction. The responses to the 5-HT doses were normalised to the tension developed in response to maximal K^+^ (0 mM NaCl, 25 mM NaHCO_3_, 4.7 mM KCl, 1.2 mM MgSO_4_.7H_2_O, 125 mM KH_2_PO_4_, 2.5 mM CaCl_2_, 2.8 mM d-glucose; Sigma-Aldrich, Gillingham, UK). Uterine vascular endothelium-dependent dilator function was also determined by measuring tension changes to cumulative increasing doses of methacholine (10^-9^ to 10^-4^ M) following pre-constriction with a sub-maximal dose of 5-HT. The methacholine dose-response curve was repeated following a 20 min pre-incubation with nitric oxide (NO) synthase inhibitor L-NAME (NG-nitro-L-arginine methyl ester hydrochloride) to determine the contribution of NO-dependent mechanisms to methacholine-mediated vasodilatation, as previously established (Herrera *et al*., 2010). Uterine vascular smooth-muscle dependent dilator function was determined by measuring tension changes to cumulative increasing doses of sodium nitroprusside (SNP; 10^-10^ to 10^-4^ M) following pre-constriction with a sub-maximal dose of 5-HT. Vessel tension changes for determining constrictor or dilator reactivity in the uterine artery were recorded with LabChart software (LabChart 6.0, Powerlab 8/30; AD Instruments, Chalgrove, UK).

### Biochemical assays

Plasma insulin levels were quantified with an ovine-specific insulin ELISA kit (Mercodia, Uppsala, Sweden). The intra-assay coefficient of variation was 3%. Total plasma cholesterol (CHOL) and triglycerides (TG) were quantified by enzymatic assays, performed by the MRC MDU Mouse Biochemistry Laboratory [MC_UU_00014/5]. Reagents were provided by Siemens Healthcare (Forchheim, Germany) and analysed on the Siemens Dimension EXL analyser. The limit of detection was 1.3 mmol/L and 0.17 mmol/L for CHOL and TG, respectively.

### Statistical analyses

All data are presented as the mean ± SEM. Statistical analyses were performed with GraphPad Prism 9.5.1 software, with *P*<0.05 considered statistically significant. Individual comparisons between CON and OB ewes were made by the Student’s *t* test for unpaired data. Diet and gestational age comparisons across longitudinal maternal samples were made using repeated measures two-way ANOVA. When a significant (*P*<0.05) interaction was present in ANOVA, the *post-hoc* Šídák’s comparison test was used to isolate significant relationships. Maternal diet, fetal sex and placentome type comparisons were made using three-way or two-way ANOVA, as appropriate. When no fetal sex effect was present (*P*>0.05 in ANOVA), male and female data were combined. In addition, all data were assessed for the impact of twinning and maternal instrumentation status via a generalised linear model (GLM) in R version 4.4.1. Groups were combined when no significant effect was present. For instance, fetal weights were included from both instrumented and non-instrumented mothers. In cases of same-sex twins, one twin was chosen at random to be included, to ensure that each pregnancy rather than fetus was the distinct biological replicate. Mixed sex-twins were not included in this study. This resulted in data available from *n*= 13 CON male (11 singleton, 2 twin), *n*= 10 CON female (9 singleton, 1 twin), *n*= 11 OB male (10 singleton, 1 twin), *n*= 10 OB female (9 singleton, 1 twin) fetuses.

## Results

### Ewe weight gain, body condition and dietary intake

Analysis of food intake in a subset of ewes showed that the OB group (*n*= 21) consumed 967±20 g of concentrated feed per day, compared to 200±0 g in the CON group (*n*= 20). Therefore, OB relative to CON ewes had a *ca.* five-fold increase in daily energy intake. While there was no difference between groups in body weight or condition score at baseline, ewes in the OB group were significantly heavier and displayed an elevated body condition score after 4 weeks of diet exposure (Fig 1A & 1B). This increased weight gain trajectory continued throughout the pre-pregnancy feeding period, such that OB ewes entered pregnancy 30% heavier than CON ewes (Fig 1A). OB ewes continued to gain more weight throughout pregnancy, such that their total percentage weight gain was 57% by 120 dGA, compared to 12% in CON ewes (Fig 1C). By post-mortem at 130 dGA, OB ewes had significantly greater adiposity, with a 115% increase in pericardial fat and a 183% increase in perirenal fat, relative to CON ewes (Fig 1D & 1E).

**Figure 1.**
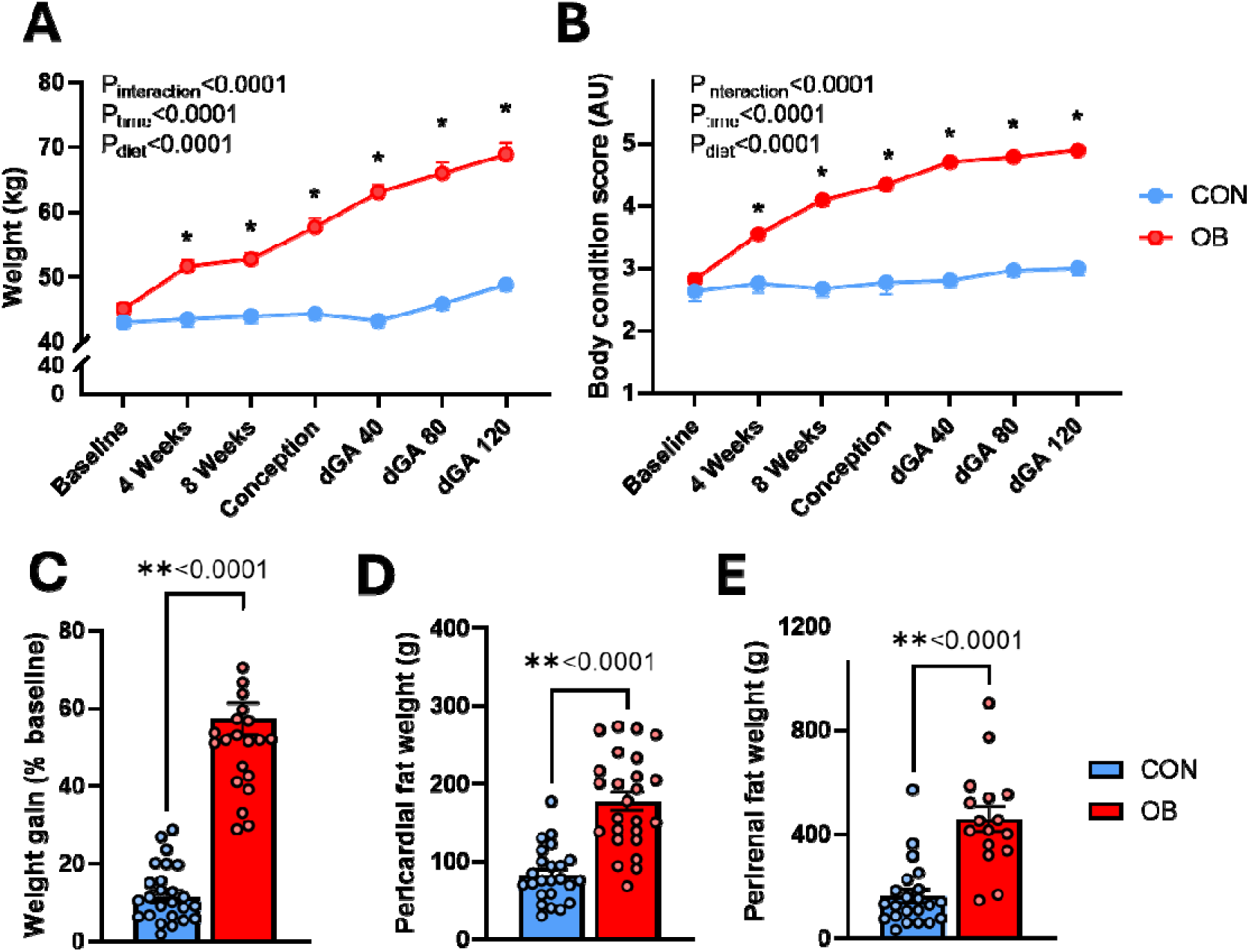
Ewe weight gain and adiposity. A) Ewe body weight and B) condition score across the pre-pregnancy, conception and pregnancy periods. C) Total maternal weight gain by 120 dGA, expressed as a percentage of baseline weight, and D) maternal pericardial and E) perirenal fat deposit weights upon post-mortem at 130 dGA. All data are the mean ± SEM in control (CON; blue) and obese (OB, red) ewes, with *n*= 18-41 ewes per diet group, per time point. \**P*<0.003 CON vs. OB comparison; Šídák’s multiple comparison test following *P*<0.0001 *Diet x Time* interaction in two-way repeated measures ANOVA. \*\**P*<0.001; unpaired *t*-test. AU; arbitrary units.

### Longitudinal in vivo metabolic phenotype

Prior to pregnancy, there was no difference in blood glucose levels between ewe groups. However, OB ewes showed a significant *Diet x Time* interaction in blood glucose levels, where OB relative to CON mothers exhibited an 18-23% elevation in blood glucose, from conception through to 120 dGA (Fig 2A). This was accompanied by a significant *Diet x Time* interaction in circulating insulin levels (Fig 2B). Across the pre-pregnancy period, the insulin AUC was not significantly different between CON and OB ewes (0.38±0.18 CON *vs*. 0.59±0.27 OB; arbitrary units (AU), Fig 2B).

**Figure 2.**
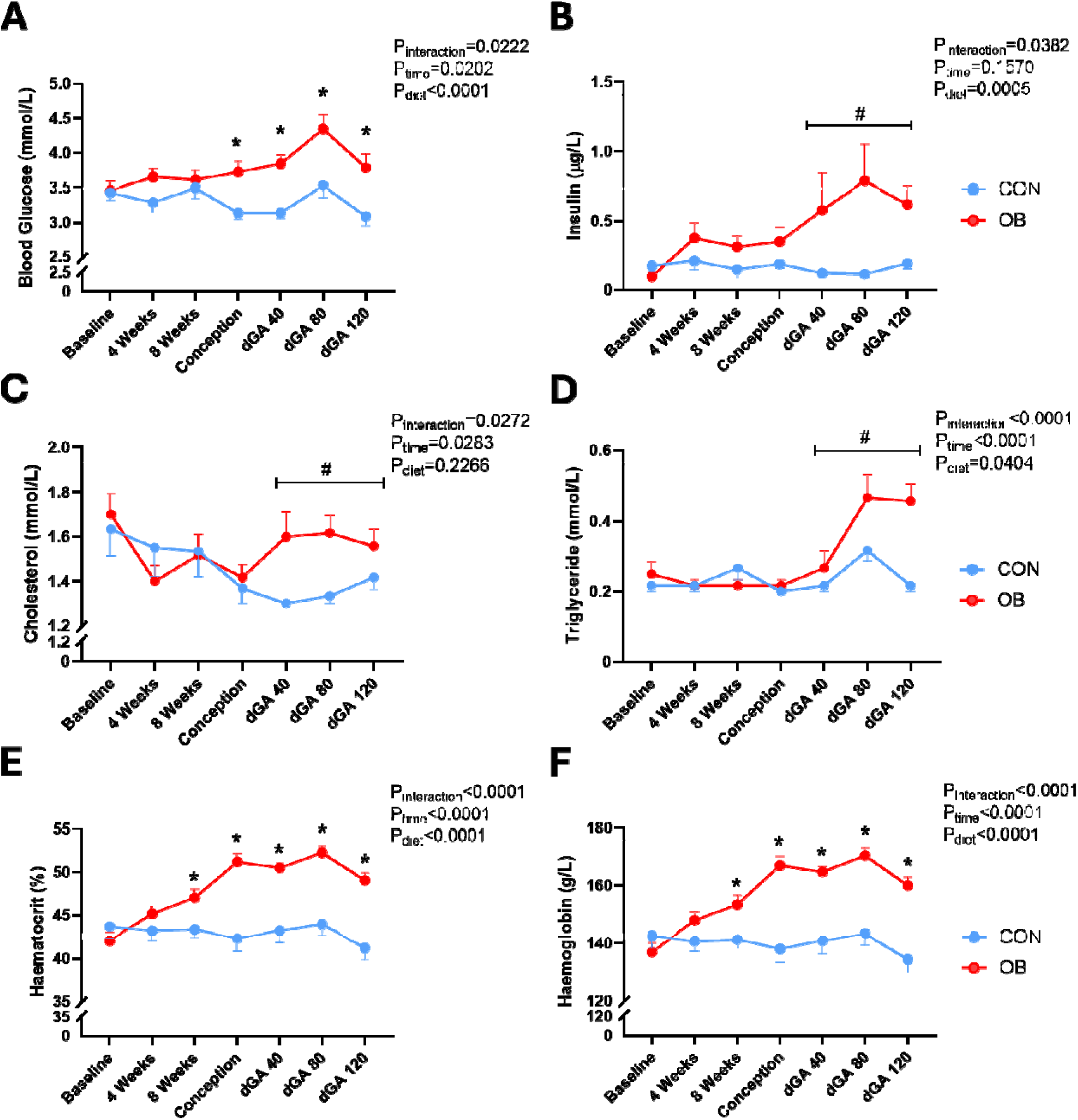
Maternal circulating metabolic markers. A) Ewe blood glucose, B) plasma insulin, C) plasma cholesterol, D) plasma triglycerides, E) haematocrit and F) haemoglobin across the pre-pregnancy, conception and pregnancy periods. All data are the mean ± SEM, with *n*= 15-28 ewes per diet group, per time point for glucose, haemoglobin and haematocrit, and *n*= 6–10 ewes per diet group, per time point for plasma markers. \**P*<0.05; CON vs. OB comparison; Šídák’s multiple comparisons test following *Diet x Time* interaction in two-way repeated measures ANOVA. #*P*<0.005; CON vs. OB AUC comparison during pregnancy; Šídák’s multiple comparisons test following *Diet x Pregnancy Status* interaction in two-way ANOVA.

However, OB relative to CON ewes showed a significantly greater insulin AUC throughout pregnancy (0.43±0.11 AU CON *vs.* 1.85±0.73 AU OB; *P*<0.001). Similarly, total plasma cholesterol and triglyceride levels displayed *Diet x Time* interactions, where circulating lipid levels were no different between ewe groups in the pre-pregnancy period. However, OB relative to CON ewes had significantly greater plasma lipid levels across pregnancy (Fig 2C & 2D). Specifically, the AUC of both cholesterol (2.70±0.09 AU CON *vs.* 3.20±0.23 AU OB; *P*=0.0013) and triglycerides (0.53±0.06 AU CON *vs.* 0.83±0.14 AU OB; *P*<0.001) was significantly elevated in OB ewes during pregnancy (Fig 2C & 2D). Further, OB relative to CON ewes, showed significantly higher haematocrit and haemoglobin levels from 8 weeks of pre-pregnancy feeding through to 120 dGA (Fig 2E & 2F).

### Maternal in vivo systemic cardiovascular function and nutrient delivery

During basal measurements, OB relative to CON ewes were hypertensive, showing a 19% elevation in mean arterial blood pressure (Fig 3A). This manifested as elevated systolic and diastolic pressure in OB ewes (Fig 3B & 3C). While basal heart rate was similar between diet groups (Fig 3D), maternal femoral arterial blood flow was elevated by 45% and femoral vascular resistance was reduced by 26% in OB relative to CON ewes (Fig 3E & 3F). There was no difference in arterial oxygen saturation between ewe groups, but OB relative to CON ewes had significantly greater haemoglobin concentration (Fig 3G & 3H). This resulted in arterial oxygen content and oxygen delivery in the femoral arterial circulation to be 18% and 74% greater in OB relative to CON ewes, respectively (Fig 3J & 3K). In contrast, despite greater blood glucose levels in venous blood sampling from conception through to 120 dGA in non-instrumented OB relative to CON ewes (Fig 2A), arterial blood glucose levels in the instrumented ewes were no different between diet groups (Fig 3K). However, glucose delivery to the femoral arterial circulation remained significantly elevated by 53% in OB relative to CON ewes (Fig 3L).

**Figure 3.**
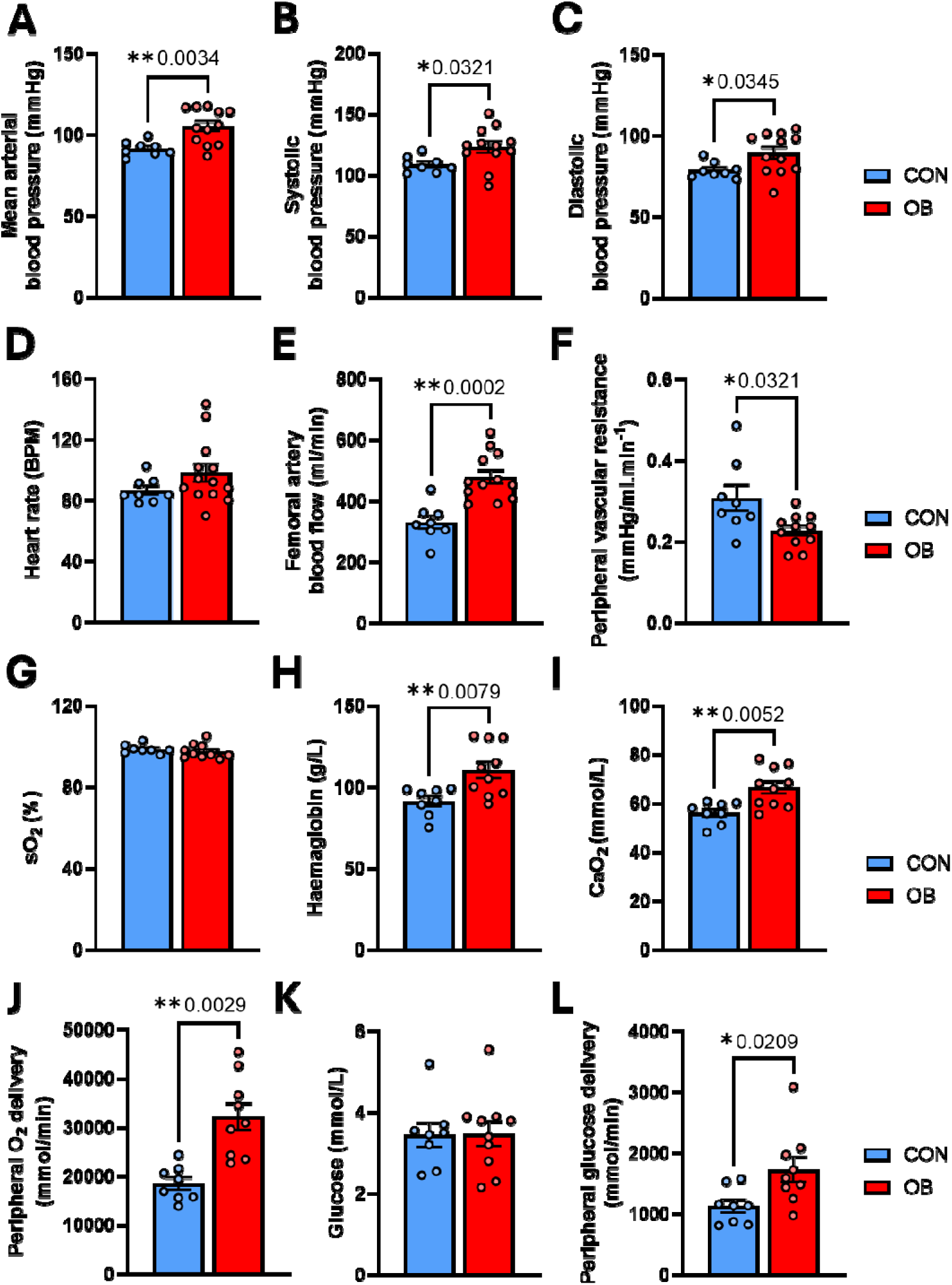
Maternal cardiovascular function in late pregnancy. Maternal A) mean blood pressure, B) systolic blood pressure, C) diastolic blood pressure, D) heart rate, E) femoral artery blood flow, F) femoral vascular resistance, G) oxygen saturation, H) haemoglobin, I) oxygen content, J) peripheral oxygen delivery, K) blood glucose and L) peripheral glucose delivery in control (CON; blue) and obese (OB, red) ewes at 120-130 dGA. All data are the mean ± SEM, with *n*= 7-12 ewes per diet group. \**P*<0.05; \*\**P*<0.01; \*\*\**P*<0.001; unpaired *t*-test.

### Maternal in vivo uterine vascular function and nutrient delivery

During basal measurements, OB relative to CON ewes had similar uterine artery blood flow, uterine vascular resistance, uterine oxygen delivery and uterine glucose delivery (Fig 4A, 4C, 4E & 4G). However, in contrast to other outcome variables, there were significant *Maternal Diet x Fetal Sex* interactions in uterine blood flow and vascular resistance, whereby the uterine artery blood flow tended to be lower and uterine vascular resistance was 43% higher in OB relative to CON mothers carrying a female fetus (Fig 4B & 4D). There was also a significant *Maternal Diet x Fetal Sex* interaction in uterine oxygen delivery, whereby OB mothers carrying a male fetus tended to have higher oxygen transport, relative to CON (Fig 4F). Uterine glucose delivery was not impacted by maternal obesity or fetal sex (Fig 4H).

**Figure 4.**
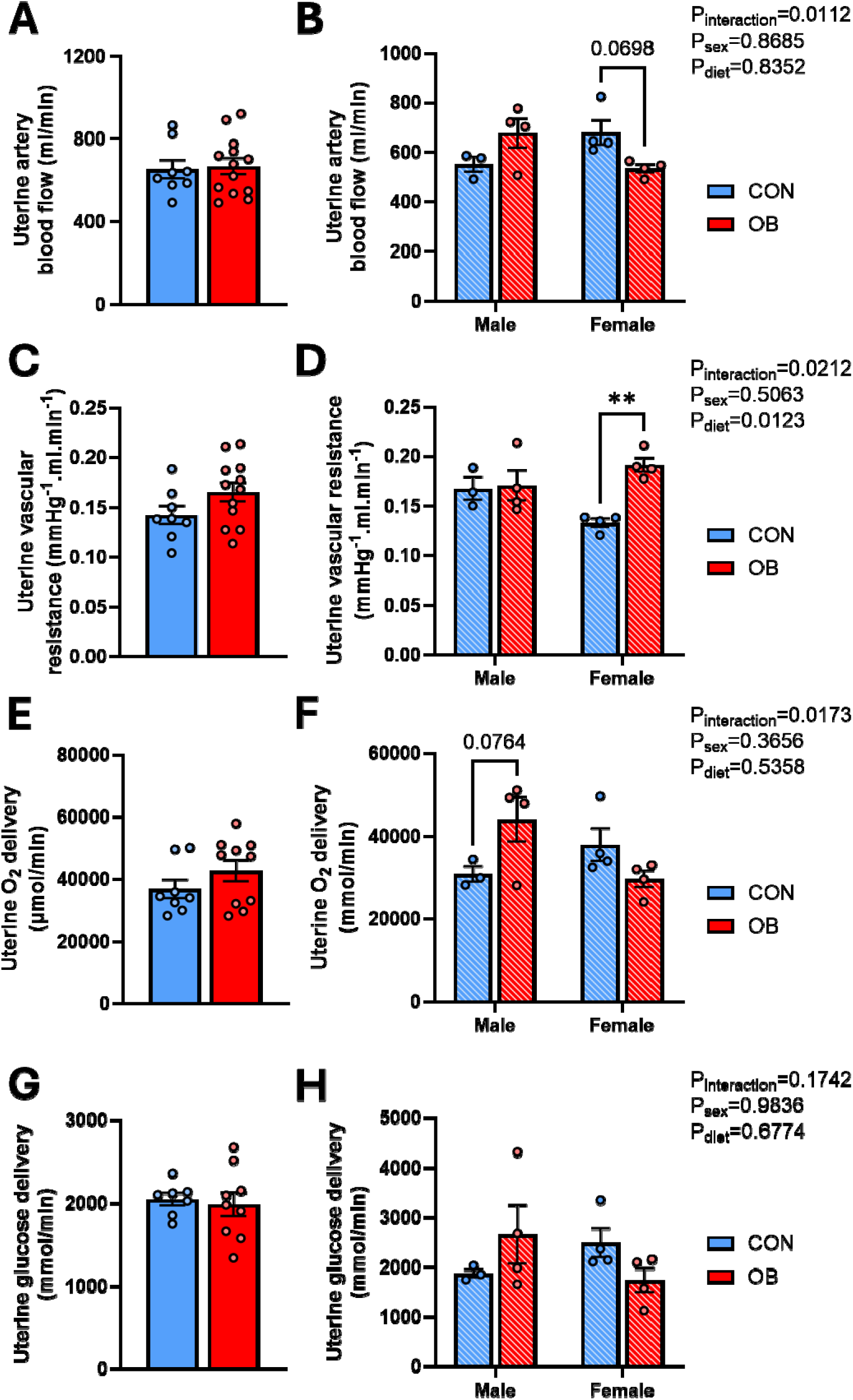
Uterine artery vascular function in late pregnancy. Uterine artery A) blood flow, B) blood flow separated by fetal sex, C) vascular resistance, D) vascular resistance separated by fetal sex, E) oxygen transport, F) oxygen transport separated by fetal sex, G) glucose transport, and H) glucose transport separated by fetal sex in control (*n*= 8; blue) and obese (*n*= 12; red) ewes at 120-130 dGA. All data are the mean ± SEM. \*\**P*<0.005; CON vs. OB comparison; Šídák’s multiple comparisons test following *Diet x Fetal Sex* interaction in two-way ANOVA.

### Maternal venous oxygenation, metabolic and electrolyte status at dGA 130

Just prior to euthanasia at 130 dGA, OB ewes showed an increase in venous pO2, but no difference in pCO2 or oxygen saturation compared to CON (Table 1). Corresponding to the longitudinal profile, OB relative to CON ewes had elevated haemoglobin and haematocrit (Table 1). OB mothers were also acidotic compared to CON, with a lower pH and a reduction in bicarbonate levels, resulting in a significant reduction in acid-base excess, relative to controls (Table 3). OB relative to CON ewes also had significantly greater values for circulating venous blood glucose, lactate, sodium, and chloride levels (Table 1).

**Table 1.** Maternal venous oxygenation, metabolic and electrolyte status at 130 dGA.

| BLOOD MARKER | CON |  | OB |  | P-Value |
| --- | --- | --- | --- | --- | --- |
|  | MEAN | SEM | MEAN | SEM |  |
| Oxygenation |  |  |  |  |  |
| pCO<sub>2</sub> (mmHg) | 41.7 | 1.17 | 41.7 | 1.66 | NS |
| pO<sub>2</sub> (mmHg) | 41.1 | 1.42 | 46.4 | 2.07 | 0.0429 |
| sO<sub>2</sub> (%) | 62.1 | 2.10 | 64.4 | 3.34 | NS |
| Haemoglobin (g/L) | 128 | 2.47 | 149 | 3.67 | <0.0001 |
| Haematocrit (%) | 39.0 | 0.78 | 45.7 | 1.13 | <0.0001 |
| Acid-Base status |  |  |  |  |  |
| pH | 7.36 | 0.01 | 7.29 | 0.02 | 0.0063 |
| Bicarbonate (mmol/L) | 22.4 | 0.70 | 18.8 | 0.80 | 0.0019 |
| Base Excess (mmol/L) | -1.75 | 0.91 | -6.64 | 1.22 | 0.0031 |
| Metabolic status |  |  |  |  |  |
| Glucose (mmol/L) | 3.66 | 0.21 | 4.43 | 0.23 | 0.0184 |
| Lactate (mmol/L) | 7.10 | 0.74 | 9.87 | 0.88 | 0.0219 |
| Electrolytes |  |  |  |  |  |
| K<sup>+</sup> (mmol/L) | 5.33 | 0.93 | 4.66 | 0.13 | NS |
| Na<sup>+</sup> (mmol/L) | 149 | 0.60 | 151 | 0.45 | 0.0116 |
| Ca<sup>2+</sup> (mmol/L) | 1.15 | 0.02 | 1.14 | 0.02 | NS |
| Cl<sup>-</sup> (mmol/L) | 108 | 0.51 | 110 | 0.49 | 0.0098 |

All data are the mean ± SEM. *P* values are derived from unpaired *t*-tests between control (CON; *n*= 18) and obese (OB; *n*= 17 -18) ewes at 130 dGA. NS; non-significant.

### Maternal, fetal and placental biometry at post-mortem

At post-mortem, at 130 dGA, OB relative to CON ewes showed greater adiposity (Table 2 and Fig 1D & 1E), and increased mass in several metabolic and endocrine organs, including an increase in heart, adrenal gland, kidney, liver, spleen, and thyroid gland weight (Table 2). When organ weights were expressed per kg of maternal body weight, relative heart rate was significantly lower, and relative pericardial and perirenal fat deposits remained significantly elevated in OB relative to CON ewes consistent with increased fat mass preferentially contributing to the increase in body weight (Table 2).

**Table 2.**
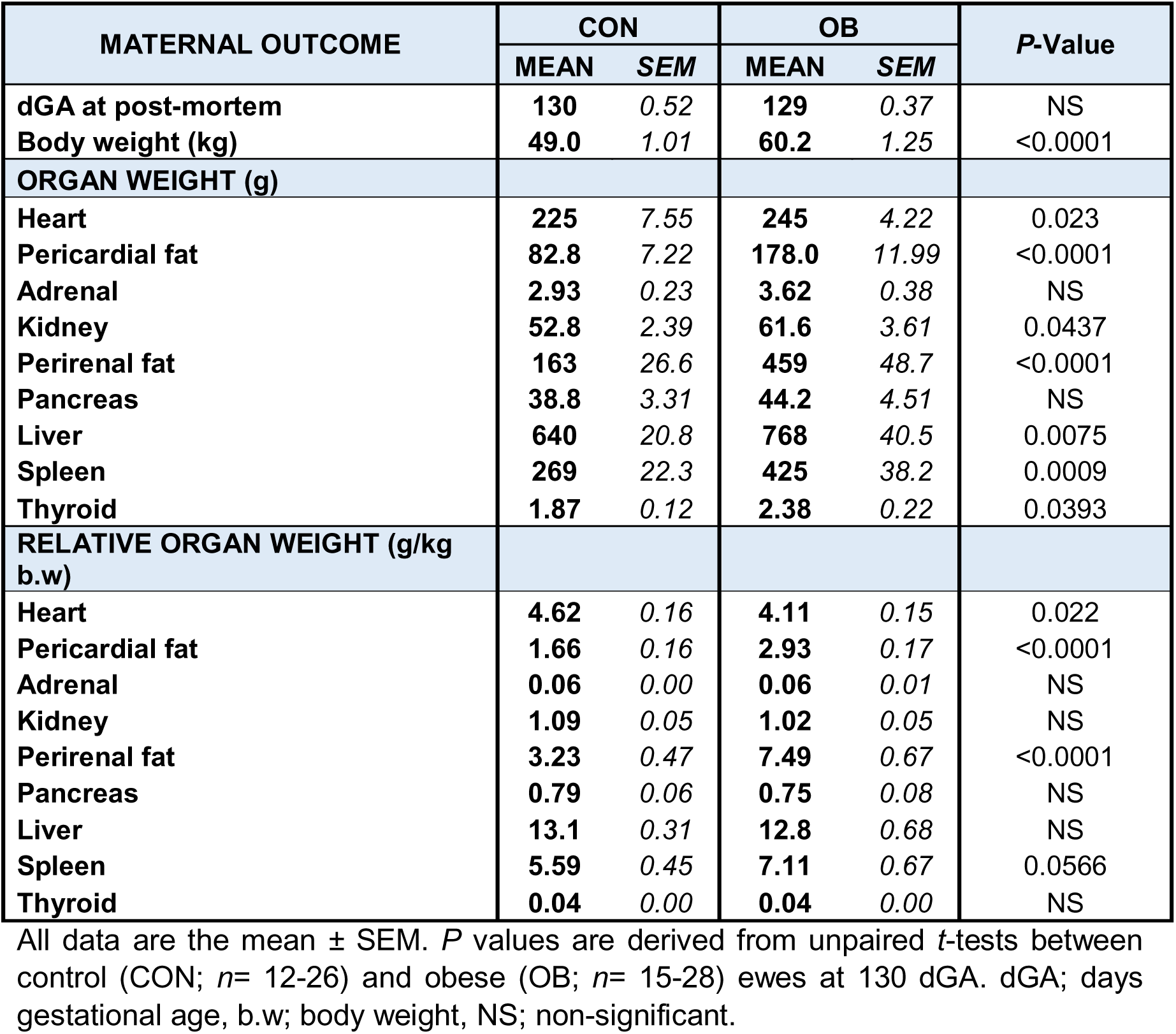
Maternal biometry at 130 dGA.

At post-mortem, at 130 dGA, fetal body weight, placental weight and placental efficiency (fetal body weight/placental weight) were similar in CON and OB groups (Fig 5A-5C). When placentomes were typed, mean overall placentome type weight but not number was significantly elevated in OB relative to CON ewes (Fig 5D & 5E). Therefore, placentomal efficiency was significantly reduced in OB relative to CON ewes (Fig 5F). At post-mortem, fetuses from OB relative to CON ewes also showed an increase in CRL, BPD, and femur length, resulting in significantly lower values for ponderal index (Table 3). Values for absolute but not relative heart, lung, kidney and perirenal fat were also significantly increased in fetuses from OB relative to CON ewes (Table 3).

**Figure 5.**
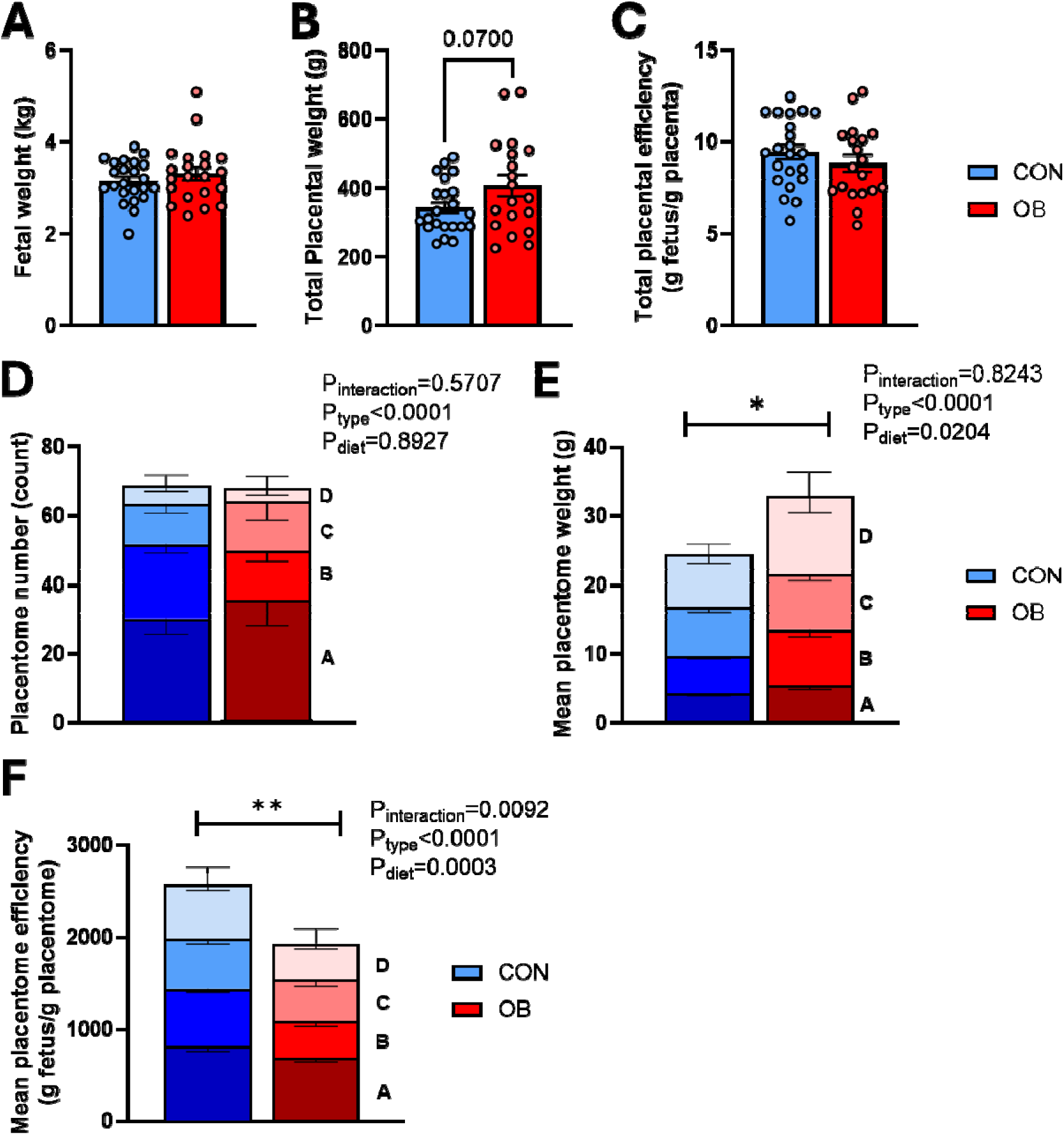
Fetal and placental phenotype at 130 dGA. A) Fetal weight, B) total placental weight, C) total placental efficiency, D) placentome number, E) mean placentome weight and F) mean placentome efficiency in control (CON; blue; *n*= 21) and obese (OB; red; *n*= 14) pregnancies at 130 dGA. All data are the mean ± SEM. No fetal sex differences were observed (*P*>0.05; two-way ANOVA), so male and female values are combined. \**P*<0.05, \*\**P*<0.001; *Diet* effect in two-way ANOVA.

**Table 3.** Fetal biometry at 130 dGA.

| FETAL OUTCOME | CON |  | OB |  | P-Value |
| --- | --- | --- | --- | --- | --- |
|  | MEAN | SEM | MEAN | SEM |  |
| dGA at post-mortem | 130 | 0.52 | 129 | 0.37 | NS |
| Body weight (kg) | 3.15 | 0.09 | 3.32 | 0.14 | NS |
| **MORPHOMETRY** |  |  |  |  |  |
| Crown rump length (cm) | 42.4 | 0.56 | 44.2 | 0.64 | 0.0329 |
| Biparietal diameter (cm) | 9.56 | 0.26 | 11.05 | 0.35 | 0.0013 |
| Abdominal circumference (cm) | 29.5 | 0.61 | 29.8 | 1.37 | NS |
| Femur length (cm) | 11.3 | 0.26 | 12.2 | 0.29 | 0.0390 |
| BMI (weight (kg)/CRL (cm)<sup>2</sup>) | 17.6 | 0.42 | 16.8 | 0.40 | NS |
| PI (weight (g) x 100) / CRL (cm)<sup>3</sup>) | 4.16 | 0.12 | 3.81 | 0.10 | 0.0337 |
| **ORGAN WEIGHT (g)** |  |  |  |  |  |
| Heart | 22.3 | 0.71 | 24.5 | 0.94 | 0.0604 |
| Brain | 41.8 | 1.03 | 44.8 | 1.10 | 0.0522 |
| Right lung | 41.7 | 1.41 | 47.9 | 2.67 | 0.0379 |
| Left lung | 28.2 | 1.01 | 32.3 | 2.09 | 0.0665 |
| Pericardial fat | 7.54 | 0.44 | 8.50 | 0.63 | NS |
| Adrenal | 0.17 | 0.01 | 0.20 | 0.01 | NS |
| Kidney | 8.51 | 0.32 | 10.38 | 0.82 | 0.0212 |
| Perirenal fat | 5.90 | 0.28 | 7.06 | 0.47 | 0.0328 |
| Pancreas | 2.36 | 0.15 | 2.72 | 0.21 | NS |
| Liver | 82.0 | 3.61 | 95.7 | 7.16 | 0.0884 |
| Spleen | 5.07 | 0.21 | 5.86 | 0.41 | 0.0705 |
| Thyroid | 0.33 | 0.02 | 0.36 | 0.03 | NS |
| **RELATIVE ORGAN WEIGHT (g/kg b.w)** |  |  |  |  |  |
| Heart | 7.11 | 0.17 | 7.51 | 0.29 | NS |
| Brain | 13.6 | 0.51 | 13.6 | 0.41 | NS |
| Right lung | 12.3 | 0.72 | 13.8 | 0.42 | NS |
| Left lung | 8.34 | 0.49 | 9.28 | 0.31 | NS |
| Pericardial fat | 2.39 | 0.12 | 2.53 | 0.12 | NS |
| Adrenal | 0.06 | 0.00 | 0.06 | 0.00 | NS |
| Kidney | 2.55 | 0.18 | 2.93 | 0.14 | NS |
| Perirenal fat | 1.84 | 0.07 | 1.99 | 0.09 | NS |
| Pancreas | 0.74 | 0.04 | 0.73 | 0.07 | NS |
| Liver | 26.6 | 1.45 | 28.6 | 1.92 | NS |
| Spleen | 1.65 | 0.08 | 1.70 | 0.09 | NS |
| Thyroid | 0.10 | 0.01 | 0.10 | 0.01 | NS |

All data are the mean ± SEM. *P* values are derived from unpaired *t*-tests between control (CON; *n*=19-23) and obese (OB; *n*=12-21) fetuses at 130 dGA. There was no significant effect of fetal sex on any outcome (*P*>0.05; two-way ANOVA), so male and female data are combined. NS; non-significant, BMI; body mass index, PI; ponderal index.

### Uterine artery vascular reactivity measured ex vivo via in vitro wire myography

Vascular smooth muscle constriction induced by serotonin (5-HT) involves Rho kinase (Fig 6A). Uterine vascular constrictor reactivity to 5-HT was similar before, but significantly attenuated following blockade with the synthetic Rho kinase inhibitor Y27632 in CON relative to OB ewes, independent of the sex of the fetus (Fig 6B & 6C). This suggests that OB ewes rely more on Rho kinase-independent mechanism to mediate the constrictor response to 5-HT in the uterine vasculature.

**Figure 6.**
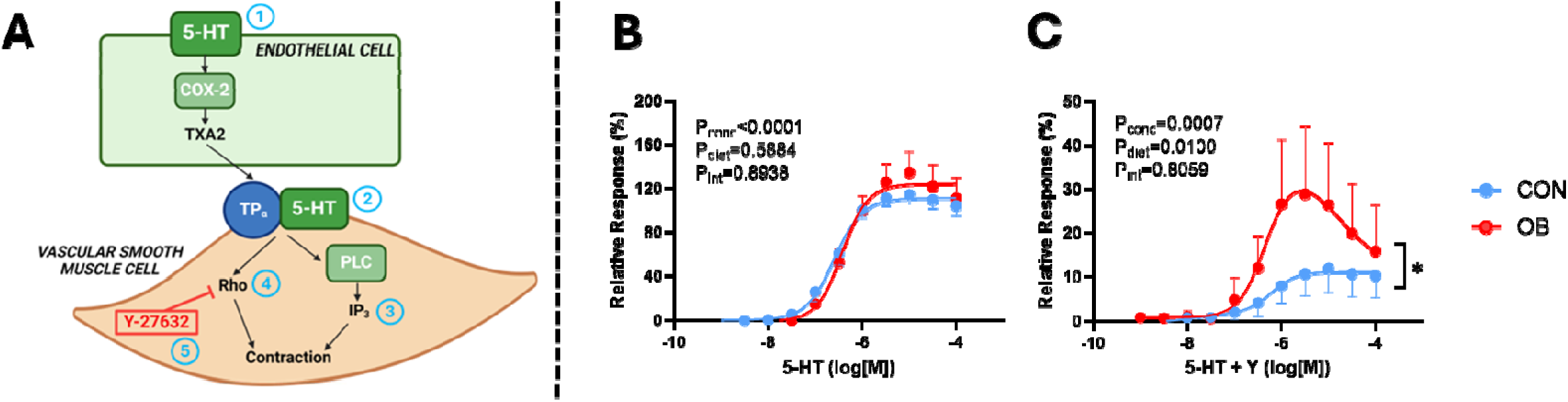
Uterine artery vascular constrictor reactivity at dGA 130. A) diagram of vasoconstrictor pathways in the uterine artery: *Serotonin (5-HT) can induce vasoconstriction via endothelium-dependent thromboxane A2 (TXA2) production in endothelial cells (1) or by direct action on vascular smooth muscle cells (2). Activation of phospholipase C (PLC) produces inositol triphosphate (IP3) (3), which releases Ca2+ from the sarcoplasmic reticulum to activate myosin light chain kinase, thereby stimulating vascular smooth muscle cell contraction. Alternatively, Rho kinase can directly phosphorylate myosin light chain and inhibit myosin light chain phosphatase activity to stimulate contraction (4). Rho kinase activity is blocked by the synthetic inhibitor, Y27632 (Y) (5).* Uterine artery concentration response curves to B) serotonin (5-HT) and C) 5-HT+ Y in control (CON; blue) and obese (OB; red) ewes at 130 dGA. All data are the mean ± SEM with *n*= 10-13 ewes per diet group. \**P*<0.05 Diet effect in two-way ANOVA.

Uterine vascular dilator reactivity to the smooth muscle-dependent agonist sodium nitroprusside (SNP; Fig 7A) was similar in OB relative to CON ewes when all ewes were included (Fig 7B). However, there was a significant M*aternal Diet x Fetal Sex* interaction (*P*=0.0039; Fig 7B), whereby ewes carrying a female, but not male fetus, displayed impaired uterine artery relaxation in response to lower doses of SNP relative to CON (Fig 7C & 7D).

**Figure 7.**
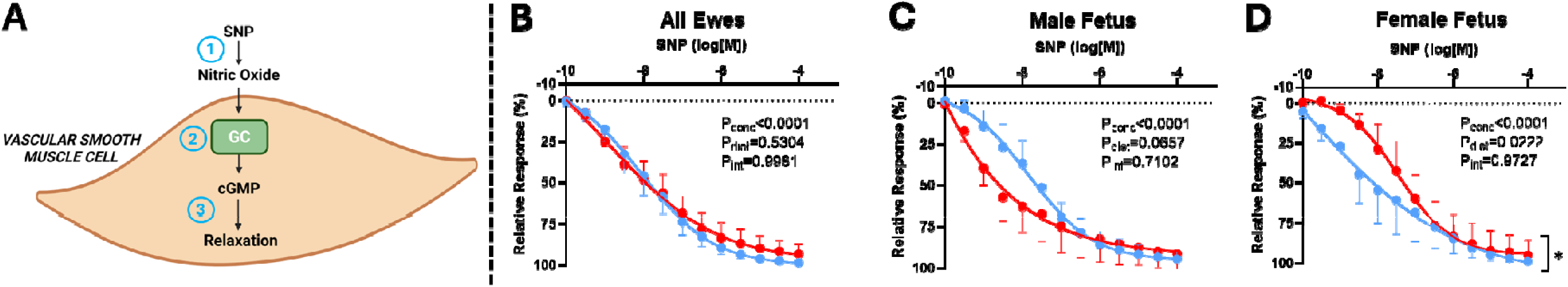
Uterine artery vascular smooth muscle-dependent dilator reactivity at dGA 130. A) diagram of vascular smooth muscle-dependent vasodilator pathways in the uterine artery: *Sodium nitroprusside (SNP) breaks down to produce nitric oxide (1). This stimulates guanylate cyclase (GC) in vascular smooth muscle cells to activate cyclic GMP (cGMP) (2). cGMP reduces intracellular calcium levels, which induces vasorelaxation (3).* Uterine artery concentration response curves to sodium SNP in A) all ewes, B) ewes carrying a male fetus, C) ewes carrying a female fetus. All data are the mean ± SEM with *n*= 10-13 ewes per diet group in combined ewe analysis, *n*= 4-7 ewes per group in separate fetal sex analysis. \**P*<0.05 Diet effect in two-way ANOVA.

Uterine artery vascular dilator reactivity to the endothelium-dependent agonist methacholine (MCh; Fig 8A) was significantly impaired when all ewes were included before but not after blockade with L-NAME in OB relative to CON ewes (Fig 8B & 8E). However, there was a significant interaction between maternal diet and fetal sex. OB ewes carrying a male had greater vasorelaxation to MCh before (Fig 8C) and following L-NAME (Fig 8F), while OB ewes carrying a female fetus were unaffected (Fig 8D & 8G). This suggests that OB ewes carrying a male fetus have greater NO-independent mechanisms mediating the uterine vasodilator response to MCh.

**Figure 8.**
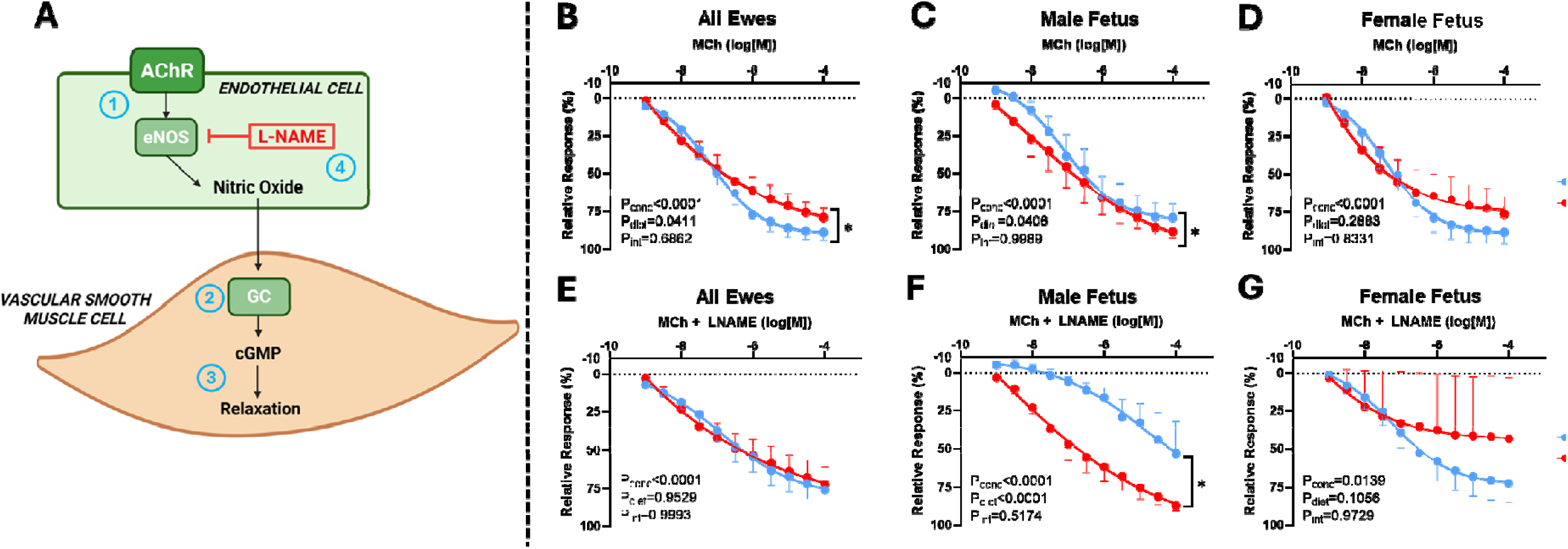
Uterine artery vascular endothelial-dependent dilator reactivity at dGA 130. A) diagram of vascular endothelial-dependent vasodilator pathways in the uterine artery: *Methacholine (MCh) binds to vascular endothelial cell acetylcholine receptors (AChR) to induce endothelial nitric oxide synthase (eNOS) (1). This produces nitric oxide, which stimulates guanylate cyclase (GC) in vascular smooth muscle cells to activate cyclic GMP (cGMP) (2). cGMP reduces intracellular calcium levels, which induces vasorelaxation (3). Administration of L-NAME directly blocks eNOS action in endothelial cells, thereby preventing MCh-induced vasodilation via this pathway (4).* Uterine artery concentration response curves to A) MCh in all ewes, B) MCh in ewes carrying a male fetus, C) MCh in ewes carrying a female fetus, D) MCh+LNAME in all ewes, E) MCh+LNAME in ewes carrying a male fetus, and F) MCh+LNAME in ewes carrying a female fetus, in control (CON; blue) and obese (OB; red) ewes at 130 dGA. All data are the mean ± SEM with *n*= 10-13 ewes per diet group in combined ewe analysis, n= 2-7 ewes per group in separate fetal sex analysis. \**P*<0.05 Diet effect in two-way ANOVA.

## Discussion

This study introduces a novel ovine model of diet-induced obesity during pregnancy, where ewes with obesity display increased adiposity, hyperglycaemia, hyperinsulinemia, hyperlipidaemia, and hypertension, constituting cardiometabolic dysfunction typical of the metabolic syndrome. Relative to lean controls, ewes with obesity also showed increased haemoglobin throughout the study period, with elevated femoral blood flow and enhanced peripheral oxygen transport near term. Maternal uterine artery function was significantly affected by obesity, showing increased vascular resistance *in vivo*. This was explained in part by a greater reactivity to Rho kinase-independent constrictor mechanisms stimulated by serotonin and reduced dilator responses to the endothelium-dependent agonist methacholine *ex vivo*. Effects on maternal uterine vasoreactivity in obese ewes varied according to the sex of the fetus, showing reduced smooth-muscle dependent vasorelaxation to sodium nitroprusside *ex vivo* in mothers with obesity carrying a female, but not male, fetus. Conversely, mothers with obesity carrying a male, but not female, fetus showed greater NO-independent mechanisms mediating the uterine vasodilator response to methacholine *ex vivo*. These cardiometabolic disturbances occurred with evidence of decreased placental efficiency and alterations in fetal growth, promoting a thin-for-length phenotype with a reduced ponderal index in fetuses of both sexes in ewes with obesity.

Ewes in the obese group gained weight steadily throughout the pre-pregnancy feeding period and entered pregnancy substantially heavier than controls. Moreover, expected metabolic disturbances associated with excess adiposity, including hyperglycaemia, hyperinsulinemia and hyperlipidaemia, developed in the obese ewes with pregnancy onset and progression. This reinforces the concept that pregnancy itself is a metabolic challenge that exacerbates underlying subclinical pathologies, as noted in cases of gestational diabetes mellitus (Catalano, 2014).

Interestingly, pregnant ewes with obesity also exhibited a striking increase in haemoglobin and haematocrit relative to controls, which started after eight weeks of obesogenic feeding and persisted throughout pregnancy. Elevated haemoglobin has been noted in recent cohorts of pregnant women with obesity relative to lean-weight women (Elmugabil *et al*., 2017; Eltayeb *et al*., 2023) and higher haemoglobin levels, particularly in the first trimester, have been positively associated with elevated BMI and GDM onset in larger-scale population studies (Wang *et al*., 2018; Sissala *et al*., 2022). The mechanisms underlying this elevated haemoglobin profile in women with obesity is unclear, but may be driven by maternal insulin status, since insulin can stimulate erythropoietin production by activating the HIF-1 signalling cascade (Treins *et al*., 2002) and/or by acting directly as a growth factor (Masuda *et al*., 1997).

The elevated maternal haemoglobin levels in obese ewes may also contribute to their hypertensive phenotype, since heightened haemoglobin levels in the first trimester have been positively associated with the onset of pregnancy-induced hypertension (Aghamohammadi *et al*., 2011; Abumohsen *et al*., 2021) and preeclampsia (Wang *et al*., 2018) in otherwise healthy women. Mechanisms linking elevated haemoglobin to hypertension may include an increase in blood viscosity, promoting enhanced peripheral vascular resistance according to Poiseuille’s Law and, thereby, an increase in cardiac afterload (Letcher *et al*., 1981; Çınar *et al*., 1999). Since diastolic arterial blood pressure reflects downstream peripheral vascular resistance, its elevation in pregnant ewes with obesity supports this contention. Additionally, ewes with obesity in our study had greater extracellular fluid (ECF) concentrations of sodium and reduced heart weight in relation to their total body weight compared to lean ewes. Greater sodium will lead to ECF volume expansion, increasing total fluid volume, which may have contributed to the raised blood pressure in pregnant ewes with obesity. A relatively smaller heart to perfuse a larger body, combined with higher blood viscosity and cardiac afterload, will contribute to higher cardiac workload in pregnant ewes with obesity. Significantly greater systolic pressure in pregnant ewes with obesity supports this contention. Increased cardiac workload, hypertension and higher haemoglobin levels are all strongly associated with long-term cardiometabolic disease risk (Mehta & Dubrey, 2009; Honigberg *et al*., 2019; Tapio *et al*., 2021), and so may perpetuate cardiometabolic dysfunction in mothers with obesity long after birth.

Peripheral vascular function was altered substantially in pregnant ewes with obesity near term, with elevated femoral blood flow, reduced femoral vascular resistance, and enhanced oxygen and glucose supply to peripheral vascular beds, relative to controls. Elevated blood flow in the peripheral vasculature has also been previously noted in pregnant women with obesity (Dutta *et al*., 2020) and may be facilitated by the high volume to low-resistance circulation profile noted in obese relative to control pregnancies (Vonck *et al*., 2019). However, sustained elevations in blood flow can induce endothelial dysfunction by the activation of shear stress pathways, leading to vascular remodelling and changes to vascular tone (Lu & Kassab, 2011).

The elevated oxygen delivery in pregnant ewes with obesity is somewhat counter-intuitive, since obesity is typically associated with adipose tissue hypoxia in non-pregnant individuals (Ye, 2009; Mirabelli *et al*., 2024), and placental hypoxia in rodent models of obese pregnancy (Fernandez-Twinn *et al*., 2017; Wallace *et al*., 2019). While the elevated haemoglobin content in ewes with obesity clearly underlies the elevated oxygen transport findings, the haemoglobin-driven hyperviscosity may also reduce perfusion at the microvascular level (Mirabelli *et al*., 2024). Therefore, there may be a disconnect between oxygen transport levels based on arterial conduit calculations, and those occurring at terminal arterial branches. Assessment of tissue-specific oxygenation levels would be required to confirm this.

Uterine artery function was altered by maternal obesity in a fetal sex-specific manner, with elevated uterine artery vascular resistance and impaired smooth muscle-dependant relaxation capacity in pregnant ewes with obesity carrying a female fetus. In healthy pregnancy, the uterine vascular bed is subject to critical remodelling, resulting in lower uterine vascular resistance towards term, which facilitates maximal oxygen and nutrient supply to the growing fetus (Osol & Mandala, 2009). Therefore, ewes with obesity carrying a female fetus appear more vulnerable to reduced uterine nutritional supply, and our *ex vivo* data suggest that this occurs through vascular smooth muscle alterations. Interestingly, obese ewes carrying a male fetus did not exhibit any significant *in vivo* uterine artery changes relative to controls. Rather, our *ex vivo* data suggest that mothers with obesity carrying a male fetus showed greater NO-independent mechanisms mediating the uterine vasodilator response to methacholine ex vivo. This may act as a compensatory mechanism to maintain appropriate uterine artery vasomotor function, which is consistent with other evidence of maternal obesity leading to recruitment of NO-independent compensatory mechanisms for vasodilatation in a mouse model of preeclampsia (Binder *et al*., 2025). Interestingly, the elevated haemoglobin levels in OB mothers may contribute to this adaptation by reducing NO bioavailability, since haemoglobin can oxidise NO to generate nitrates and methaemoglobin (Gow *et al*., 1999; Keszler *et al*., 2008). We have not assessed nitrate levels in the present study, however further studies into NO metabolism in this model appear warranted considering these uterine artery adaptations.

Previous studies have found that women carrying a male fetus exhibit a higher uterine artery Doppler pulsatility index (PI) compared to those carrying a female fetus (Broere-Brown *et al*., 2016; Teulings *et al*., 2020). Teulings and colleagues (2020) further demonstrated that maternal obesity increases uterine PI, but this occurred independently of fetal sex effects (Teulings *et al*., 2020). To our knowledge, our study is the first to assess the interaction between maternal obesity and fetal sex on uterine artery function. Our model found significant *Maternal Diet x Fetal Sex* interactions in most uterine artery outcomes. Therefore, further investigation into the importance of maternal-fetal crosstalk in obese pregnancy is merited, and our study highlights specific mechanistic pathways to target. While there was no difference in fetal weight between diet groups at 130 dGA, fetuses exposed to maternal obesity showed a reduced ponderal index. Fetal growth outcomes in obese pregnancy are complex and somewhat heterogenic between diverse populations and models, and different studies have reported fetal overgrowth, undergrowth or no change to fetal weight with maternal obesity (Crew *et al*., 2016; Lewandowska, 2021; Langley-Evans *et al*., 2022). Our model reproducing decreased placental efficiency with fetuses which are thin-for-their-length is interesting, as it is this phenotype in offspring of complicated pregnancy that has been linked with future cardiovascular risk (Eriksson *et al*., 2001).

In summary, this study presents a robust model of maternal obesity in pregnancy that exhibits cardiometabolic dysfunction near term and recapitulates key aspects of the human phenotype, including increased adiposity, hyperglycaemia, hyperinsulinemia, hyperlipidaemia, and hypertension in the mother. Further, we provide mechanistic insights into fetal sex-specific uterine vascular bed dysfunction that may have significant implications for the maternal cardiovascular adaptation to pregnancy, and the long-term maternal and offspring cardiovascular health after pregnancy complicated by obesity. The fetal sex-specific changes to maternal uterine artery function in obese ewes also highlight the importance of fetal-to-maternal communication in regulating maternal vascular function during pregnancy.

## Author contributions

Study concept and funding acquisition: DAG, SEO, MPM. Model optimisation and study design: DAG, RCC, SGF. *In vivo* experiments and tissue generation: RCC, ALKC, YN, SGF, CLRC, SHD, DAG. Lab work and biochemical assays: RCC, ALKC. Data analyses and results interpretation: RCC, ALKC, DAG. Manuscript draft: RCC, DAG. Manuscript revision and approval: RCC, ALKC, YN, SGF, CLRC, SHD, MPM, SEO, DAG.

## Data availability statement

Data will be made available by the authors upon reasonable request.

## References

(2012). Improving Bioscience Research Reporting: The ARRIVE Guidelines for Reporting Animal Research. Veterinary Clinical Pathology 41, 27–31.

Abumohsen H, Bustami B, Almusleh A, Yasin O, Farhoud A, Safarini O, Thabaleh A, Sukhon M, Nazzal Z & Damiri B. (2021). The Association Between High Hemoglobin Levels and Pregnancy Complications, Gestational Diabetes and Hypertension, Among Palestinian Women. Cureus 13, e18840.

Aghamohammadi A, Zafari M & Tofighi M. (2011). High maternal hemoglobin concentration in first trimester as risk factor for pregnancy induced hypertension. Caspian J Intern Med 2, 194–197.

Allison BJ, Brain KL, Niu Y, Kane AD, Herrera EA, Thakor AS, Botting KJ, Cross CM, Itani N, Shaw CJ, Skeffington KL, Beck C & Giussani DA. (2020). Altered Cardiovascular Defense to Hypotensive Stress in the Chronically Hypoxic Fetus. Hypertension 76, 1195–1207.

Allison BJ, Brain KL, Niu Y, Kane AD, Herrera EA, Thakor AS, Botting KJ, Cross CM, Itani N, Skeffington KL, Beck C & Giussani DA. (2016). Fetal in vivo continuous cardiovascular function during chronic hypoxia. The Journal of Physiology 594, 1247–1264.

Barry JS & Anthony RV. (2008). The pregnant sheep as a model for human pregnancy. Theriogenology 69, 55–67.

Binder NK, de Alwis N, Fato BR, Beard S, Mangwiro YTM, Kadife E, Brownfoot F & Hannan NJ. (2025). Investigating the Impact of Maternal Obesity on Disease Severity in a Mouse Model of Preeclampsia. Nutrients 17.

Bonds DR, Crosby LO, Cheek TG, Hägerdal M, Gutsche BB & Gabbe SG. (1986). Estimation of human fetal-placental unit metabolic rate by application of the Bohr principle. Journal of developmental physiology 8, 49–54.

Brain KL, Allison BJ, Niu Y, Cross CM, Itani N, Kane AD, Herrera EA, Skeffington KL, Botting KJ & Giussani DA. (2019). Intervention against hypertension in the next generation programmed by developmental hypoxia. PLOS Biology 17, e2006552.

Broere-Brown ZA, Schalekamp-Timmermans S, Hofman A, Jaddoe V & Steegers E. (2016). Fetal sex dependency of maternal vascular adaptation to pregnancy: a prospective population-based cohort study. Bjog 123, 1087–1095.

Catalano PM. (2014). Trying to understand gestational diabetes. Diabetic Medicine 31, 273–281.

Çınar Y, Demir G, Paç M & Çınar AB. (1999). Effect of hematocrit on blood pressure via hyperviscosity. American Journal of Hypertension 12, 739–743.

Cochrane ALK, Murphy MP, Ozanne SE & Giussani DA. (2024). Pregnancy in obese women and mechanisms of increased cardiovascular risk in offspring. Eur Heart J 45, 5127–5145.

Crew RC, Mark PJ, Clarke MW & Waddell BJ. (2016). Obesity disrupts the rhythmic profiles of maternal and fetal progesterone in rat pregnancy. Biology of Reproduction 95, 55, 51–10.

Delaey C, Boussery K & Van de Voorde J. (2002). Contractility Studies on Isolated Bovine Choroidal Small Arteries: Determination of the Active and Passive Wall Tension–Internal Circumference Relation. Experimental Eye Research 75, 243–248.

Dutta EH, Burns RN, Pacheco LD, Marrs CC, Koutrouvelis A & Koutrouvelis GLO. (2020). Lower Extremity Blood Flow Velocity in Obese versus Nonobese Pregnant Women. Am J Perinatol 37, 384–389.

Elmugabil A, Rayis DA, Abdelmageed RE, Adam I & Gasim GI. (2017). High level of hemoglobin, white blood cells and obesity among Sudanese women in early pregnancy: a cross-sectional study. Future Sci OA 3, Fso182.

Eltayeb R, Binsaleh NK, Alsaif G, Ali RM, Alyahyawi AR & Adam I. (2023). Hemoglobin Levels, Anemia, and Their Associations with Body Mass Index among Pregnant Women in Hail Maternity Hospital, Saudi Arabia: A Cross-Sectional Study. Nutrients 15.

Eriksson JG, Forsén T, Tuomilehto J, Osmond C & Barker DJ. (2001). Early growth and coronary heart disease in later life: longitudinal study. Bmj 322, 949–953.

Fernandez-Twinn DS, Gascoin G, Musial B, Carr S, Duque-Guimaraes D, Blackmore HL, Alfaradhi MZ, Loche E, Sferruzzi-Perri AN, Fowden AL & Ozanne SE. (2017). Exercise rescues obese mothers’ insulin sensitivity, placental hypoxia and male offspring insulin sensitivity. Scientific Reports 7, 44650.

Gardner DS, Giussani DA & Fowden AL. (2003). Hindlimb glucose and lactate metabolism during umbilical cord compression and acute hypoxemia in the late-gestation ovine fetus. Am J Physiol Regul Integr Comp Physiol 284, R954–964.

Gow AJ, Luchsinger BP, Pawloski JR, Singel DJ & Stamler JS. (1999). The oxyhemoglobin reaction of nitric oxide. Proc Natl Acad Sci U S A 96, 9027–9032.

Grundy D. (2015). Principles and standards for reporting animal experiments in The Journal of Physiology and Experimental Physiology. J Physiol 593, 2547–2549.

Herrera EA, Verkerk MM, Derks JB & Giussani DA. (2010). Antioxidant treatment alters peripheral vascular dysfunction induced by postnatal glucocorticoid therapy in rats. PLoS One 5, e9250.

Honigberg MC, Zekavat SM, Aragam K, Klarin D, Bhatt DL, Scott NS, Peloso GM & Natarajan P. (2019). Long-Term Cardiovascular Risk in Women With Hypertension During Pregnancy. J Am Coll Cardiol 74, 2743–2754.

Kent L, McGirr M & Eastwood KA. (2024). Global trends in prevalence of maternal overweight and obesity: A systematic review and meta-analysis of routinely collected data retrospective cohorts. Int J Popul Data Sci 9, 2401.

Kenyon PR, Maloney SK & Blache D. (2014). Review of sheep body condition score in relation to production characteristics. New Zealand Journal of Agricultural Research 57, 38–64.

Keszler A, Piknova B, Schechter AN & Hogg N. (2008). The reaction between nitrite and oxyhemoglobin: a mechanistic study. J Biol Chem 283, 9615–9622.

Langley-Evans SC, Pearce J & Ellis S. (2022). Overweight, obesity and excessive weight gain in pregnancy as risk factors for adverse pregnancy outcomes: A narrative review. J Hum Nutr Diet 35, 250–264.

Letcher RL, Chien S, Pickering TG, Sealey JE & Laragh JH. (1981). Direct relationship between blood pressure and blood viscosity in normal and hypertensive subjects: Role of fibrinogen and concentration. The American Journal of Medicine 70, 1195–1202.

Lewandowska M. (2021). Maternal Obesity and Risk of Low Birth Weight, Fetal Growth Restriction, and Macrosomia: Multiple Analyses. Nutrients 13.

Lu D & Kassab GS. (2011). Role of shear stress and stretch in vascular mechanobiology. J R Soc Interface 8, 1379–1385.

Ma Y, Zhu MJ, Uthlaut AB, Nijland MJ, Nathanielsz PW, Hess BW & Ford SP. (2011). Upregulation of growth signaling and nutrient transporters in cotyledons of early to mid-gestational nutrient restricted ewes. Placenta 32, 255–263.

Masuda S, Chikuma M & Sasaki R. (1997). Insulin-like growth factors and insulin stimulate erythropoietin production in primary cultured astrocytes. Brain Research 746, 63–70.

Mehta PA & Dubrey SW. (2009). High output heart failure. QJM: An International Journal of Medicine 102, 235–241.

Mirabelli M, Misiti R, Sicilia L, Brunetti FS, Chiefari E, Brunetti A & Foti DP. (2024). Hypoxia in Human Obesity: New Insights from Inflammation towards Insulin Resistance-A Narrative Review. Int J Mol Sci 25.

Morrison JL, Berry MJ, Botting KJ, Darby JRT, Frasch MG, Gatford KL, Giussani DA, Gray CL, Harding R, Herrera EA, Kemp MW, Lock MC, McMillen IC, Moss TJ, Musk GC, Oliver MH, Regnault TRH, Roberts CT, Soo JY & Tellam RL. (2018). Improving pregnancy outcomes in humans through studies in sheep. American Journal of Physiology-Regulatory, Integrative and Comparative Physiology 315, R1123–R1153.

Osol G & Mandala M. (2009). Maternal uterine vascular remodeling during pregnancy. Physiology (Bethesda) 24, 58–71.

Owens JA, Falconer J & Robinson JS. (1987). Effect of restriction of placental growth on oxygen delivery to and consumption by the pregnant uterus and fetus. Journal of developmental physiology 9, 137–150.

Parikh NI, Gonzalez JM, Anderson CAM, Judd SE, Rexrode KM, Hlatky MA, Gunderson EP, Stuart JJ, Vaidya D, On behalf of the American Heart Association Council on E, Prevention, Council on Arteriosclerosis T, Vascular B, Council on C, Stroke N & the Stroke C. (2021). Adverse Pregnancy Outcomes and Cardiovascular Disease Risk: Unique Opportunities for Cardiovascular Disease Prevention in Women: A Scientific Statement From the American Heart Association. Circulation 143, e902–e916.

Regnault TR, de Vrijer B, Galan HL, Wilkening RB, Battaglia FC & Meschia G. (2013). Umbilical uptakes and transplacental concentration ratios of amino acids in severe fetal growth restriction. Pediatr Res 73, 602–611.

Schon SB, Cabre HE & Redman LM. (2024). The impact of obesity on reproductive health and metabolism in reproductive-age females. Fertility and Sterility 122, 194–203.

Sissala N, Mustaniemi S, Kajantie E, Vääräsmäki M & Koivunen P. (2022). Higher hemoglobin levels are an independent risk factor for gestational diabetes. Scientific Reports 12, 1686.

Tannenbaum J & Bennett BT. (2015). Russell and Burch’s 3Rs then and now: the need for clarity in definition and purpose. J Am Assoc Lab Anim Sci 54, 120–132.

Tapio J, Vähänikkilä H, Kesäniemi YA, Ukkola O & Koivunen P. (2021). Higher hemoglobin levels are an independent risk factor for adverse metabolism and higher mortality in a 20-year follow-up. Scientific Reports 11, 19936.

Täufer Cederlöf E, Lundgren M, Lindahl B & Christersson C. (2022). Pregnancy Complications and Risk of Cardiovascular Disease Later in Life: A Nationwide Cohort Study. J Am Heart Assoc 11, e023079.

Teulings NEWD, Wood AM, Sovio U, Ozanne SE, Smith GCS & Aiken CE. (2020). Independent influences of maternal obesity and fetal sex on maternal cardiovascular adaptation to pregnancy: a prospective cohort study. International Journal of Obesity 44, 2246–2255.

Tong W, Allison BJ, Brain KL, Patey OV, Niu Y, Botting KJ, Ford SG, Garrud TA, Wooding PFB, Shaw CJ, Lyu Q, Zhang L, Ma J, Cindrova-Davies T, Yung HW, Burton GJ & Giussani DA. (2022). Chronic Hypoxia in Ovine Pregnancy Recapitulates Physiological and Molecular Markers of Preeclampsia in the Mother, Placenta, and Offspring. Hypertension 79, 1525–1535.

Treins C, Giorgetti-Peraldi S, Murdaca J, Semenza GL & Van Obberghen E. (2002). Insulin Stimulates Hypoxia-inducible Factor 1 through a Phosphatidylinositol 3-Kinase/Target of Rapamycin-dependent Signaling Pathway*. Journal of Biological Chemistry 277, 27975–27981.

Vatnick I, Schoknecht PA, Darrigrand R & Bell AW. (1991). Growth and metabolism of the placenta after unilateral fetectomy in twin pregnant ewes. Journal of developmental physiology 15, 351–356.

Vervoort D, Wang R, Li G, Filbey L, Maduka O, Brewer LC, Mamas MA, Bahit MC, Ahmed SB & Van Spall HGC. (2024). Addressing the Global Burden of Cardiovascular Disease in Women: JACC State-of-the-Art Review. Journal of the American College of Cardiology 83, 2690–2707.

Vonck S, Lanssens D, Staelens AS, Tomsin K, Oben J, Bruckers L & Gyselaers W. (2019). Obesity in pregnancy causes a volume overload in third trimester. European Journal of Clinical Investigation 49, e13173.

Wallace JG, Bellissimo CJ, Yeo E, Fei Xia Y, Petrik JJ, Surette MG, Bowdish DME & Sloboda DM. (2019). Obesity during pregnancy results in maternal intestinal inflammation, placental hypoxia, and alters fetal glucose metabolism at mid-gestation. Scientific Reports 9, 17621.

Wang C, Lin L, Su R, Zhu W, Wei Y, Yan J, Feng H, Li B, Li S & Yang H. (2018). Hemoglobin levels during the first trimester of pregnancy are associated with the risk of gestational diabetes mellitus, pre-eclampsia and preterm birth in Chinese women: a retrospective study. BMC Pregnancy and Childbirth 18, 263.

Wilkening RB, Molina RD & Meschia G. (1988). Placental oxygen transport in sheep with different hemoglobin types. Am J Physiol 254, R585–589.

Ye J. (2009). Emerging role of adipose tissue hypoxia in obesity and insulin resistance. Int J Obes (Lond) 33, 54–66.

